# EFFECTS OF TOLL-LIKE RECEPTOR 3 – DEPENDENT IMMUNE ACTIVATION IN MICE ARE SEX- AND TISSUE- SPECIFIC: IMPLICATIONS FOR ALCOHOL USE DISORDER

**DOI:** 10.64898/2026.04.21.719957

**Authors:** P. Antwi-Adjei, S. Shanmugam, B. Kisby, I. Ponomarev

## Abstract

**Background:** Alcohol use disorder (AUD) is linked to increased neuroinflammation. Alcohol (ethanol) may activate toll-like receptors, which leads to the release of inflammatory molecules that could influence AUD-related behaviors, such as increased alcohol intake. Activation of toll-like receptor 3 (TLR3) by Polyinosinic:polycytidylic acid (Poly(I:C) or PIC) is associated with escalation of alcohol consumption in male, but not female F1 hybrid mice from reciprocal crosses between FVB/NJ (FVB) and C57BL/6J (B6) strains. Little is known about the underlying mechanisms of these sex-specific behavioral effects. In this study, we investigated the effects of TLR3 activation by PIC on temporal profiles of several pro- and anti-inflammatory molecules in the blood and brain of FVB/B6 F1 hybrid male and female mice at multiple time points. We hypothesized that TLR3 – dependent immune profiles would differ between males and females, which may, at least in part, explain the observed differences in drinking behavior.

**Methods:** Male and female FVB/B6 F1 hybrid alcohol-naive mice were injected intraperitoneally with PIC (10 mg/kg) or saline. Blood and perfused brain tissues from the prefrontal cortex (PFC) and striatum were collected at 6-, 24-, and 48-hours post-injection. The expressions of *Ccl2, Ccl5, Tnf, Il-6, Il-1β, Ifng, Ifnb1*, and *Mmp9* genes were analyzed using qPCR. Protein levels of a subset of these molecules and IL-17r/a, IL-4, and IL-10 were measured in striatal samples from the same animals using ELISA.

**Results:** Activation of TLR3 by PIC triggered time-dependent, sex- and tissue-specific responses in immune genes and their proteins. PIC induced a time-dependent increase in expression of majority of the genes peaking at the 6 hr time point. Temporal immune profiles for pro-inflammatory chemokines, *Ccl2* and *Ccl5* differed between males and females in the PFC and striatum, suggesting possible sex-specific effects of these molecules on behavior. Protein levels of CCL2, CCL5, and IL-6 increased in the striatum of both sexes and correlated strongly with gene expression, with females showing somewhat higher protein fold changes. MMP-9, a key regulator of blood–brain barrier (BBB) permeability and synaptic plasticity, showed an increase in protein levels, but not mRNA levels in striatum. This pattern suggests altered blood–brain barrier (BBB) permeability, although this would require further investigation.

**Conclusion:** Our results revealed distinct TLR3-dependent immune gene and protein expression profiles in blood and brain between males and females and suggested different roles for these molecules in regulating alcohol consumption. We identified CCL2, CCL5 and MMP-9 as target molecules for investigating sex-specific behavior in the immune modulation of alcohol consumption.

## 1. INTRODUCTION

Alcohol Use Disorder (AUD) is a globally prevalent chronic brain disorder affecting millions of people. This behavior can have significant health and socio-economic implications for both the patient and society [1]. AUD development may be driven by a complex interaction between genetic/neurobiological, environmental, and psychological factors [2-4]. Chronic neuroinflammation is considered a key neurobiological factor due to its association with AUD-related behaviors like alcohol craving and negative emotional states. [5, 6]. Chronic alcohol use induces inflammation in the brain by interactions with neurons and glial cells, leading to the production of pro-inflammatory cytokines. The persistence of pro-inflammatory cytokines in the CNS may disrupt neurotransmitter systems, brain reward pathways, and affect the balance between excitatory and inhibitory neurotransmission, as well as synaptic plasticity. These alterations can occur in brain regions critical for the regulation of AUD-related behaviors, such as the prefrontal cortex (PFC) and striatum, which may contribute to excessive alcohol consumption.

Toll-like receptors (TLRs) are Pattern Recognition Receptors (PRRs) found on innate immune cells and neurons. These receptors detect ligands such as PAMPs (Pathogen-Associated Molecular Patterns) from bacteria and viruses, e.g., dsRNA and lipopolysaccharides, and DAMPs (Damage-Associated Molecular Patterns) from endogenous sources, e.g., HMGB1. More importantly, Toll-like receptor 3 can be activated by endogenous dsRNA released from damaged cells during alcohol exposure. This alcohol-induced activation of these receptors promotes neuroinflammation, contributing to the pathophysiology of Alcohol Use Disorder. Specifically, TLRs 2, 3, 4, and 7 [7-9] are found significantly upregulated in the post-mortem brains of alcoholics compared to non-alcoholics. Alcohol consumption increases TLR3/TRIF-dependent signaling in the prefrontal cortex of C57BL/6J mice, and the magnitude of this signaling positively correlates with alcohol intake [10]. In addition, direct TLR3 activation using polyinosinic:polycytidylic acid (Poly(I:C); PIC) augmented pro-inflammatory cytokine expression in both peripheral blood and brain tissue in the presence of alcohol [11] and increased alcohol consumption in male C57BL/6J mice [12]. This sex- and strain-specific behavior is also seen in male F1 hybrid mice from reciprocal crosses between FVB/NJ (FVB) and C57BL/6J (B6) mouse strains [13]. The male F1 hybrids drink more ethanol after repeated injections of PIC, whereas females show no change or decrease in ethanol consumption [13]. Also, the activation of TLR3 by Poly(I:C) produces a rapid innate immune response in males compared to female mice, which may have a delayed and prolonged response [14]. In addition, there are significant sex differences in neuroimmune genes expressed at the transcriptome level in brain of AUD murine models [15]. These suggest that specific functions of discrete immune genes may influence sex-dependent AUD-related behaviors.

To better understand how sex influences TLR3-mediated immune responses, we analyzed the temporal immune profiles of male and female FVB/B6 F1 hybrid mice after TLR3 activation with PIC. We hypothesized that the expression of specific cytokines and chemokines would differ between males and females at both mRNA and protein levels, potentially explaining differences in drinking behavior. We measured gene expression in blood and in two key brain regions (PFC and striatum), which allowed us to evaluate the relationships between peripheral and CNS inflammation. We focused on striatum to evaluate protein levels of specific cytokines and chemokines and investigate mRNA-protein correlations. Understanding sex-dependent immune responses after TLR3 activation could improve our understanding of how sex-specific treatment strategies might be developed for AUD-related behaviors, such as excessive alcohol intake.

## 2. MATERIALS AND METHODS

### 2.1 Animals and housing

Ethanol-naïve male C57BL/6J and female FVB/NJ mice were obtained from The Jackson Laboratory at 7 weeks of age and allowed a 1-week habituation period in the breeding colony before mating. FVB/B6 F1 hybrids were produced by crossing FVB females with C57BL/6J males as previously described [16-18]. Offspring were weaned at postnatal day 21 and transferred to the experimental room at least 1 week before testing. Animals were single-housed and maintained under controlled temperature and humidity on a 12:12 h light/dark cycle (lights on at 06:00 h), with ad libitum access to food and water. Mice were 8 weeks old at the time of Poly(I:C) treatment. All experimental procedures were approved by the Texas Tech University Health Sciences Center Institutional Animal Care and Use Committee and were conducted in accordance with the NIH Guide for the Care and Use of Laboratory Animals (IACUC protocol #19009).

### 2.2 Grouping and Poly(I:C) injections

Prior to injection, FVB/B6 F1 mice were counterbalanced by body weight, littermates, treatment, and sex (males, *n* = 33; females, *n* = 34). Within each sex and treatment group, tissue collection was conducted at defined intervals following Poly(I:C) or saline administration (6, 24, or 48 h post-injection). Tissues obtained from saline-treated animals (n = 8 males; n = 8 females) at the respective time points were combined to generate a unified control group for each sex and designated as time point 0 for all subsequent analyses. All mice received an intraperitoneal injection of Poly(I:C) (10 mg/kg) or saline, and tissues were harvested at the assigned time points.

### 2.3 Brain and blood Tissue isolation

For blood collection, cardiac puncture was performed to obtain 250-500 microliters of blood per mouse using sterile 1 mL syringes. Immediately following blood collection, mice were perfused with phosphate-buffered saline (PBS) while cardiac activity was maintained to remove residual blood from the brain. Blood samples were transferred directly into 2 mL microcentrifuge tubes pre-filled with TRIzol LS reagent to prevent clotting and initiate RNA stabilization and were kept on ice until further processing.

Following perfusion, brains were rapidly extracted, rinsed in PBS, and embedded in cryomolds containing optimal cutting temperature (OCT) compound (Sakura Finetek USA Inc., Torrance, CA, USA). Brains were fully submerged and oriented for coronal sectioning in cryomolds. The molds were then slowly cooled to solidification in a pre-chilled isopentane bath surrounded by dry ice. Once solidified, samples were stored at −80 °C until further use.

### 2.4 Brain slicing and punching

The OCT blocks containing the brains were carefully removed from the cryomolds and fixed onto a cryostat specimen holder using an additional layer of OCT compound for adhesion. This assembly was allowed to cool and harden in a cryostat (Leica CM 1950, Leica Biosystems, IL 60010 USA) set to a chamber temperature of -15°C before being mounted for slicing. The brains were then sliced into 300-micron-thick coronal sections. These sections were collected onto pre-cleaned RNase-treated glass slides and appropriately labeled. For each brain region (PFC, striatum), equal numbers of punches were obtained from both hemispheres using a 1.5 mm punch tool and the coordinates from the mouse brain atlas. The punches were stored in 1.5 ml DNA Lobind Eppendorf tubes at -80°C until RNA isolation.

### 2.5 Blood and Brain RNA Isolation and RT-qPCR Gene Expression

High-quality RNA was isolated from brain punches and blood samples of 67 mice. Blood RNA was extracted using TRIzol LS reagent, as described by [19], according to the manufacturer’s protocol. RNA from the PFC and striatum brain punches was extracted using the MagMAX™-96 for Microarrays Total RNA Isolation Kit (Thermo Fisher Scientific Baltics UAB, Vilnius, Lithuania). The quality of RNA from each tissue was assessed with a Nanodrop spectrophotometer (Thermo Fisher Scientific, Madison, WI, USA). The total RNA from blood and brain punches was converted to cDNA using a High-Capacity cDNA Reverse Transcription Kit (Thermo Fisher Scientific Baltics UAB, Vilnius, Lithuania). Target gene probes for *Ccl2, Ccl5, Tnf, Il6, Il1β, Mmp9, Ifng, and Ifnb1* (Thermo Fisher Scientific, TaqMan® Gene Expression Assays) were used to detect and quantify these cytokine genes with a thermal cycler (BIO RAD CFX384 Real-Time PCR System). CT values of target genes (e.g., *Ccl2, Ccl5*) were normalized to *Malat1* levels in the same samples (brain and blood). The 2–ΔΔCT method was used to calculate relative gene expression in Poly(I:C)-treated samples compared with the control (saline)

**Table 1:**
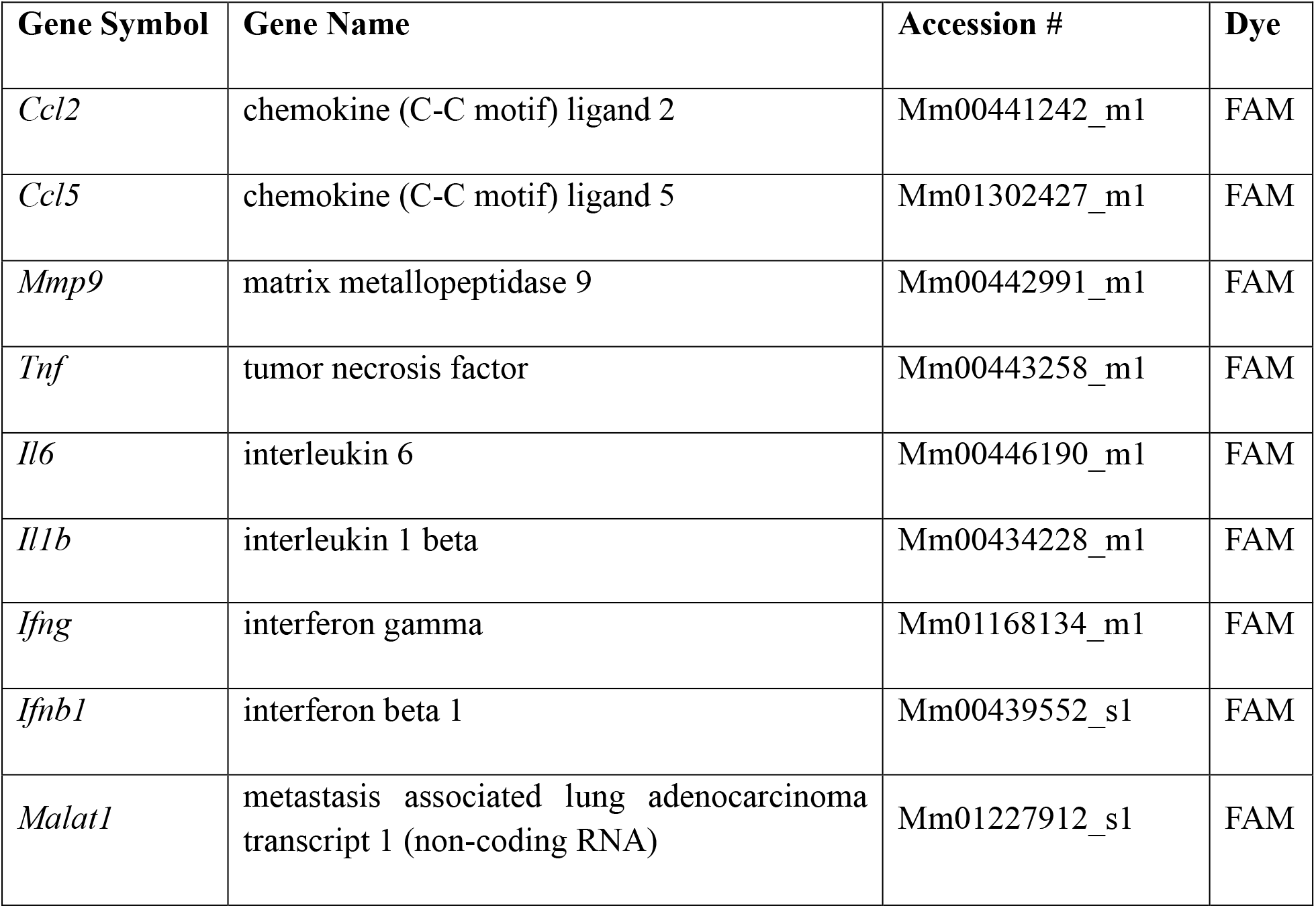
List of genes and their TaqMan probes used.

### 2.6 ELISA

To measure brain’s cytokine and chemokine protein levels, we randomly selected 40 mice for striatum punches based on sex and group (5 males and 5 females per timepoint after PIC injection, i.e., saline as 0 hours, 6 hours, 24 hours, 48 hours). We used the Meso Scale Discovery (MSD) U-PLEX kit (96-well plate), which allowed us to customize immunoassays and detect multiple cytokines and chemokines in a single well. We measured the protein levels of chemokines (CCL2, CCL5), pro-inflammatory cytokines (TNF, IL-6, IL-1β, IL-17A), anti-inflammatory cytokines (IL-4, IL-10), interferons (IFN-γ), and metalloproteinases (MMP-9). The striatum tissue was homogenized in MSD lysis buffer, and proteins were extracted by centrifugation. The extracts were normalized to a protein concentration of 3 µg/µL using lysis buffer. A 2-fold dilution of this sample mixture was prepared with an assay diluent (60 µL normalized sample + 60 µL assay diluent). Fifty microliters of each sample mixture, buffer, and freshly prepared standards were aliquoted into the 96-well plate from the kit. The plate was read with a MESO QuickPlex SQ120, and data were analyzed using the MSD Discovery Workbench software v. 4.0 (MSD, Rockville, MD, USA). The lower limit of detection (LLOD) varied by analyte, with the following levels (pg/mL): IFN-γ: 0.16; IL-1β: 3.1; IL-4: 0.56; IL-6: 4.8; IL-10: 3.8; IL-17a: 0.3; TNF-α: 1.3; MMP-9: 49; MCP-1/CCL2: 1.4; RANTES/CCL5: 0.72. Values below the LLOD were replaced with 0 pg/mL in all analyses and figures. Protein concentration levels over time were normalized to the control prior to statistical analysis.

### 2.7 Statistical analysis

Statistical analyses were performed using GraphPad Prism 11 (GraphPad Software, San Diego, CA, USA). Data are presented as mean ± SEM unless otherwise specified, and statistical significance was set at p < 0.05. Quantitative PCR (qPCR) data were analyzed using the comparative Ct method. Raw cycle threshold (Ct) values for target genes were normalized to the reference gene (*Malat1*) to obtain ΔCt values (ΔCt = Ct_target − Ct_reference). Relative gene expression was then calculated using the ΔΔCt method by normalizing each sample to the mean ΔCt of the control group.

Statistical analyses were performed on ΔΔCt values using two-way ANOVA with sex and time as factors, including their interaction (sex × time), followed by Sidak’s post hoc tests where appropriate. For visualization, gene expression data are presented as fold change values calculated as 2^(-ΔΔCt). Protein levels were normalized to control values and analyzed using the same two-way ANOVA framework (sex, time, and sex × time interaction), followed by Sidak’s post hoc tests where appropriate. Protein data are presented as normalized values. Pearson correlation analysis (α = 0.05) was used to assess associations between gene expression (ΔΔCt) and normalized protein levels. Outliers were identified using Grubbs’ test (α = 0.05)

## 3. RESULTS

### 3.1 Experiment 1: Gene expression data

Details of statistical analysis are provided in Supplementary materials.

**Poly(I:C) Elicits Sex-Specific Temporal Changes in Proinflammatory, Interferon, and *Mmp9* Signaling in Blood, PFC, and Striatum (Figure 1a)**.

**Figure 1:**
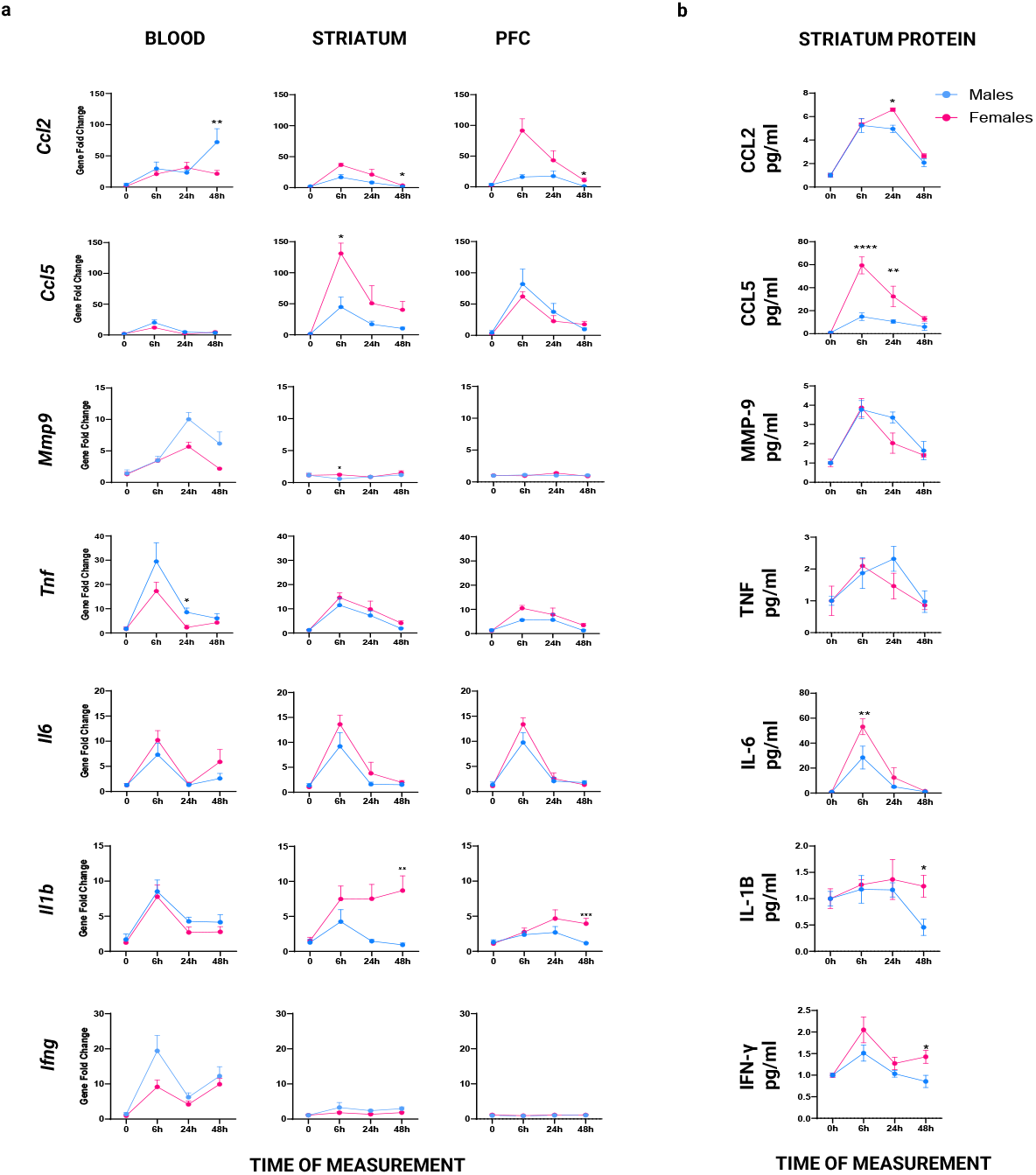
Sex-specific expression of blood, striatum, and PFC immune genes (a) and striatum immune proteins (b) at 0, 6, 24, and 48 hours after Poly(I:C) injection in FVB/B6 F1 hybrid mice.

#### 3.1.1 Blood qPCR Results

Poly(I:C) administration induced robust time-dependent increases in pro-inflammatory gene expression in blood. Males generally exhibited higher gene expression than females. *Ccl2* expression showed significant main effects of sex and time, with males displaying markedly higher levels and a pronounced peak at 48 hours. Post hoc analyses confirmed that expression was significantly elevated in males at this time point; however, no significant sex × time interaction was observed. *Ccl5* and *Il6* both exhibited significant time-dependent increases in expression. Although *Ccl5* showed a higher fold change in males and *Il6* in females, neither gene demonstrated a significant main effect of sex or a significant interaction.

*Tnf* expression showed significant main effects of sex and time, with males exhibiting higher transcript levels. Post hoc analyses indicated that this sex difference was significant at the 24-hour time point. In contrast, *Il1b* expression showed a significant main effect of time only, with no significant effects of sex or interaction. *Ifng* and *Ifnb1* both demonstrated significant time-dependent increases, with a trend toward higher expression in males; however, no significant sex or interaction effects were observed. A similar pattern was observed for *Mmp9*, which showed a significant main effect of time without significant sex differences. In summary, the overall pattern of gene expression indicates that males mount a more robust peripheral immune response than females, as reflected by both stronger trends and greater statistical significance.

#### 3.1.2 PFC qPCR Results

In the PFC, Poly(I:C) administration induced robust time-dependent increases in pro-inflammatory gene expression in the prefrontal cortex (PFC) of both sexes, with selective sex-dependent effects across genes. *Ccl2* expression exhibited significant main effects of sex and time. Post hoc analyses revealed a significant sex difference at 48 hours, with females showing higher expression. No significant sex × time interaction was observed.

*Ccl5* and *Il6* both demonstrated strong and significant time-dependent increases in expression. Although *Ccl5* exhibited a large fold change in males (close to 100-fold) and *Il6* in females (close to 80-fold), neither gene showed significant main effects of sex or interaction effects. *Tnf* expression did not show significant main effects of sex or time, nor a significant interaction even though it was more elevated in females. *Il1b* expression showed a significant main effect of sex and a significant sex × time interaction. Post hoc analyses indicated that females exhibited significantly higher expression at 48 hours. *Ifng* showed no significant main effects or interaction. In contrast, *Ifnb1* exhibited a significant sex × time interaction, driven by higher expression in females at 48 hours. Finally, *Mmp9* showed no significant main effects of sex or time, and no significant interaction effect. In conclusion, the overall pattern indicates that, aside from *Ccl5*, females exhibit greater gene expression responses in the PFC following Poly(I:C) treatment, supporting a stronger neuroimmune activation compared to males.

#### 3.1.3 Striatum qPCR Results

Poly(I:C) administration elicited time-dependent increases in immune gene expression in the striatum, with responses generally more pronounced in females. *Ccl2* expression showed a significant main effect of time and was significantly elevated in females at 48 hours; however, no significant sex × time interaction was observed. *Ccl5* exhibited significant main effects of both sex and time, with females displaying higher expression at 6 hours, corresponding to the peak response. No significant interaction effect was detected. *Il6* and *Tnf* both showed significant main effects of time only, with no significant effects of sex or interaction. *Il1b* expression demonstrated significant main effects of sex and time, with females exhibiting higher expression at 48 hours. *Ifng* showed a significant main effect of time only, with no significant effects of sex or interaction. In contrast, *Ifnb1* exhibited a significant main effect of sex, with females showing higher expression at 6 hours, but no significant interaction effect. *Mmp9* showed no significant main effects of time or interaction; however, a significant sex difference was observed at 6 hours, with females exhibiting slightly higher expression. In summary, the overall pattern of gene expression indicates that females mount a stronger and more sustained striatal immune response to Poly(I:C), as reflected by greater expression across multiple pro-inflammatory and interferon-related genes.

### 3.2 Experiment 2: Protein levels data

Details of statistical analysis are provided in Supplementary materials.

**Poly(I:C) increased striatal pro-inflammatory cytokine protein levels in both sexes, with greater fold increases in females, whereas males exhibited higher MMP-9 induction (Figure 2)**.

**Figure 2:**
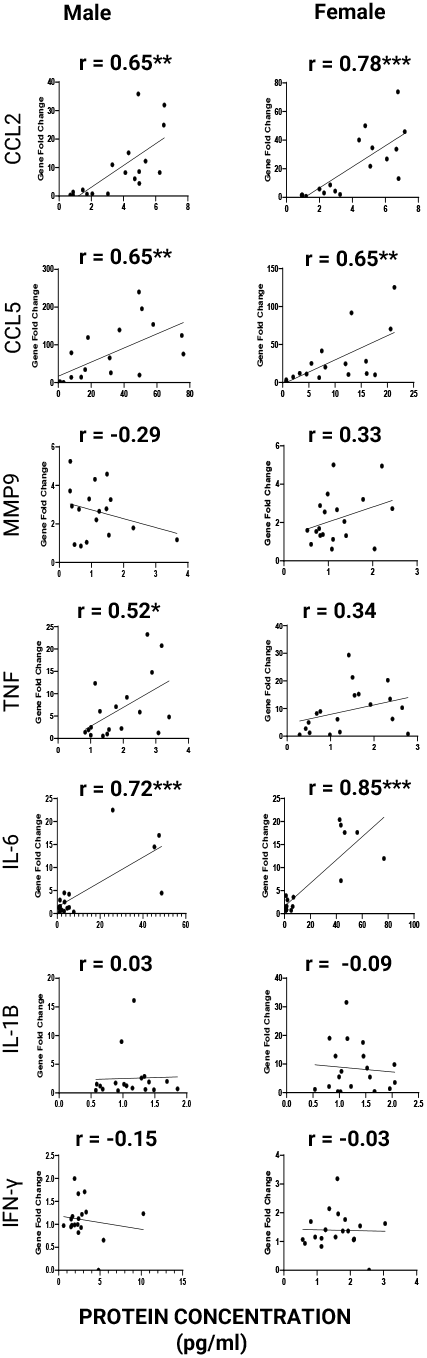
Correlation between striatal gene expression fold change and protein concentration in male and female FVB/B6 F1 hybrid mice after PIC injection.

#### 3.2.1 Striatum ELISA Result

Injecting Poly(I:C) produced time-dependent increases in several pro-inflammatory cytokine, chemokine, and interferon protein in the striatum. Except *Mmp-9*, which showed a trend toward higher protein levels in males, most analytes were elevated to a greater extent in females, although the magnitude and statistical significance varied across markers. *Ccl2* gene expression showed a significant main effect of time and sex as it was significantly elevated in females at 24 hours after Poly(I:C) injection; however, no significant sex × time interaction was observed. *Ccl5* exhibited robust main effects of both sex and time, as well as a significant sex × time interaction. Post hoc analyses revealed marked sex differences at both 6 and 24 hours, with females displaying higher levels. *Il-6* demonstrated significant main effects of sex and time, with females showing elevated levels at 6 hours. No significant interaction effect was detected.

*Tnf* expression showed a significant main effect of time only, with no significant effects of sex or sex × time interaction. In contrast, *Il-1β* did not exhibit significant main effects of sex or time or interaction. *Ifng* displayed significant main effects of both sex and time, with females exhibiting higher levels at 48 hours; however, no significant interaction effect was observed. *Mmp-9* showed a significant main effect of time and a strong trend toward a sex difference at 24 hours (p = 0.0501), with males exhibiting higher levels. Finally, the gene expression for *Il-4, Il-17* and *Il-10* showed no significant main effects of sex or time, and no significant interaction effects were observed. Taken together, the pattern of protein expression demonstrates a predominantly female-biased neuroimmune response in the striatum following Poly(I:C) administration, with consistent elevations across multiple inflammatory mediators, in contrast to a male-biased trend observed for *Mmp-9*.

### 3.3 Transcript-protein correlation

**Distinct Sex-Specific Transcript–Protein Correlation Profiles Reveal Differential Post-Transcriptional Regulation of Neuroimmune Mediators (Figure 3)**

**Figure 3:**
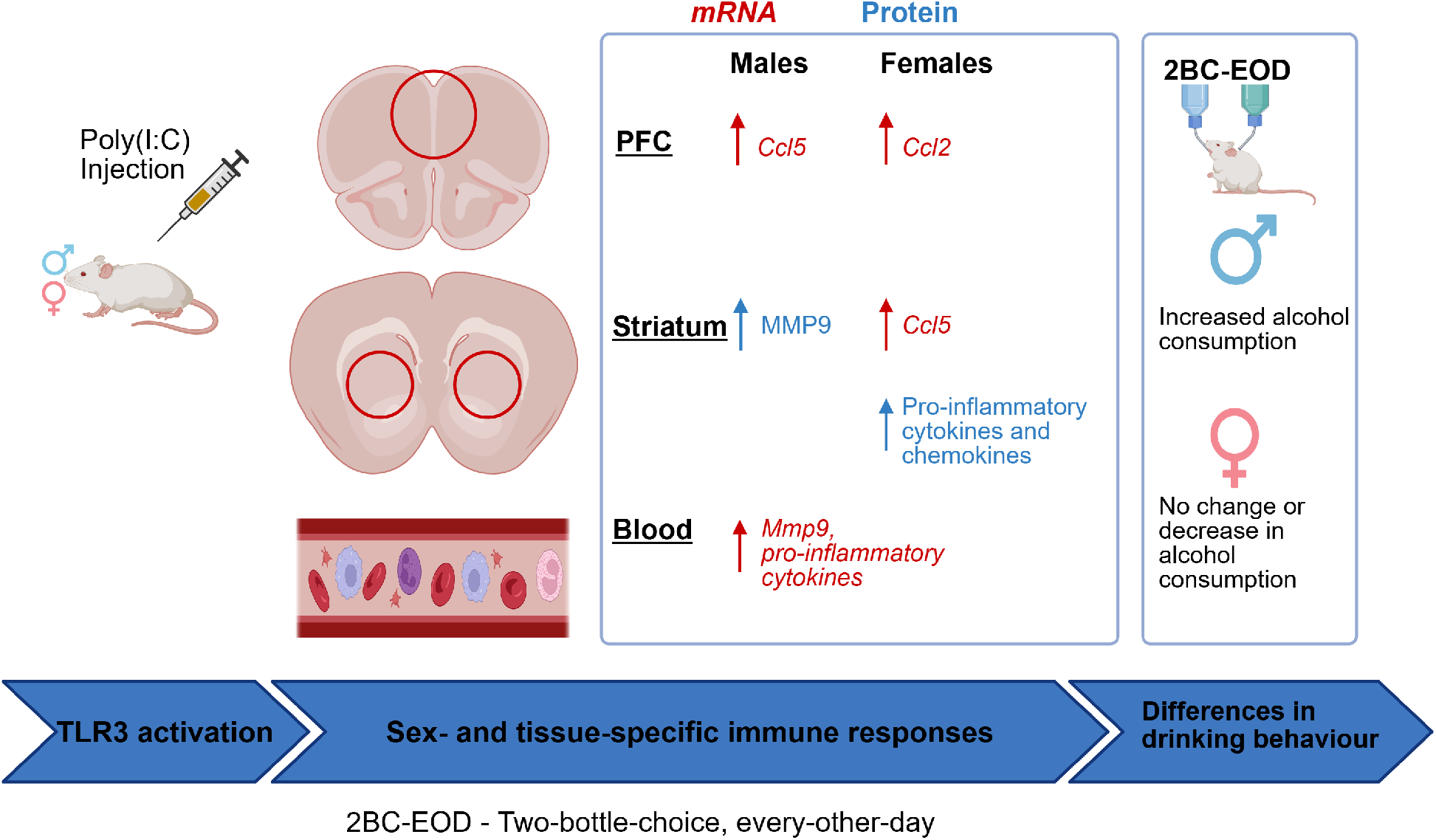
Graphical abstract showing the sex-dependent effects of TLR3 activation on immune profiles in brain and blood of FVB/B6 F1 hybrid mice.

Using a subset of mice selected for ELISA (n = 40, i.e., 20 males, 20 females), we compared normalized protein concentrations with ΔΔCT-derived gene expression changes in the striatum of the same animals. Transcript–protein correlation analyses revealed clear sex-specific regulatory patterns. Pro-inflammatory chemokines (CCL2, CCL5) and the cytokine IL-6 showed strong positive associations between gene expression and protein levels in both sexes. In contrast, TNF and IL-1β showed weaker correlations overall, although males consistently displayed relatively stronger associations than females. For IFN-γ, both males exhibited a very weak negative correlation. MMP-9 showed a weak negative correlation in males compared with a weak positive correlation in females, suggesting possible sex differences in post-transcriptional regulation. It is also possible that MMP-9 protein levels in the male striatum may not be primarily driven by local gene expression but could instead reflect contributions from peripheral sources, such as circulating blood-derived MMP-9. Collectively, transcript–protein relationships revealed sex-dependent patterns, with strong agreement for several cytokines but a distinct discrepancy for MMP-9 in males, suggesting a potential peripheral contribution to its striatal protein levels.

## 4. DISCUSSION

Persistent neuroinflammation is strongly associated with increased alcohol craving and consumption. In preclinical murine models, repeated administration of TLR3 agonist Poly(I:C) in specific mouse strains induces sustained neuroimmune activation and alters alcohol drinking behavior in a sex-dependent manner. We propose that sex-dependent differences in neuroimmune signaling pathways may partially explain variations in this AUD-related behavior. Our previous study examined the effects of TLR3-mediated immune escalation of ethanol intake and preference in FVB/NJ x C57BL/6J F1 hybrid mice [13]. We expanded on this study by examining sex differences at the molecular level in tissue-specific immune responses following TLR3 activation in FVB/B6 F1 hybrid mice. We observed sex differences in the temporal expression of pro-inflammatory cytokines, chemokines, and other immune-related genes in both blood and brain over 48 hours. We verified the expression of key cytokines at the protein level in the striatum, a brain region essential for regulating reward, habit formation, and binge drinking. Our results show distinct sex-dependent immune profiles, with CCL5 and MMP-9 identified as promising targets for future studies focusing on immune-mediated excessive alcohol intake.

Following TLR3 activation, females generally showed stronger and more prolonged brain cytokine and chemokine responses, while males exhibited more pronounced blood inflammatory gene expression. Specifically, *Tnf* gene fold change levels rose sharply in male blood but remained moderately high across all examined tissues in females. Elevated circulating TNF-α is linked to BBB dysfunction and alcohol dependence [20, 21], and this may indicate a peripheral–central signaling pathway for TLR3-driven neuroinflammation. IL-6 transcripts were strongly upregulated in both sexes, with females showing greater protein levels in the striatum, suggesting the involvement of astrocytes in promoting local inflammation [22, 23]. Elevated brain IL-6 is implicated in alcohol-related neurobiological changes, such as synaptic dysfunction [24-26] and neurobehavioral changes like craving [27]. IL-1β showed sex differences, with significantly higher transcript and protein levels in the female striatum and higher transcript levels in the PFC that persisted for 48 hours. There is multiple evidence showing that IL-1 signaling influences GABAergic transmission and ethanol consumption [28-32].

Chemokine expression also followed sex- and region-specific patterns. Females showed elevated *Ccl2* levels in the PFC, while males exhibited delayed increases in the blood. We also observed high fold changes in *Ccl5* mRNA in both the male PFC. How this may influence craving, compulsive behavior, and excessive drinking in males requires further investigation, as it will shed light on the differences in drinking behavior. In females, both CCL5 protein and gene expression levels were significantly elevated in the striatum, suggesting regionally distinct roles. These fold change patterns for *Ccl2* and *Ccl5* are similar to those already published[12, 14]. In the brain, *Ccl5* mRNA is expressed in discrete cortical and hippocampal layers, as well as in tyrosine hydroxylase–positive neurons of the ventral tegmental area (VTA), indicating expression in dopaminergic cells of the mesolimbic reward pathway [33]. The striatum, particularly the ventral striatum, which houses the nucleus accumbens, receives dense dopaminergic input from VTA neurons and forms the core of the mesolimbic reward circuit. Our observation of elevated CCL5 within the striatum suggests a role in modulating dopaminergic signaling and synaptic plasticity in reward-related pathways. There is evidence that in rats, escalating morphine use results in elevated levels of CCL5 protein and mRNA within the cortex and striatum, and this correlates with compulsive morphine use. Elevated CCL5 in the PFC and striatum modulates microglial responses, contributing to compulsive morphine use [34]. Prolonged elevation of CCL5 may also desensitize the mu-opioid receptor via cross-talk with CCR5 on neurons and glia. This can reduce reward signaling, promote anhedonia during withdrawal, and enhance neuroinflammation, collectively driving increased drug intake and reinforcing addiction-like behaviors [35].

Interferon signaling also diverged by sex, with higher transcripts in male blood and higher transcripts and protein in female brain, reflecting sex-dependent neuroimmune responses relevant to ethanol-induced TLR3 activation. Although interferons are produced in response to ethanol interacting with neurons and astrocytes [36, 37] their roles in AUD-related behaviors remain largely understudied. Elevated pro-inflammatory signaling in female brains suggests the induction of a robust sickness-like state that may suppress alcohol consumption. Increased levels of CCL2 [38], along with IL-6, IL-1β and TNF-α [39-41], are key mediators of neuroinflammation-driven sickness behavior through their effects on microglial activation and central cytokine signaling. This inflammatory profile likely reduced motivation and reward-seeking, contributing to decreased or unchanged drinking patterns in females. In contrast, males exhibit relatively lower brain cytokine expression. High *Ccl5* in the male PFC, may influence craving, compulsive behavior, and excessive drinking in the absence of sickness induction. Consequently, males may experience less sickness-related suppression and instead undergo neuroimmune changes that allow continued or escalated alcohol consumption.

MMP-9 regulation further highlighted the complexity of temporal- and tissue-specific responses. Interestingly, although *Mmp9* transcripts were elevated in male blood, there was a decreased striatal transcript expression and an increased striatal protein level. This likely reflects tissue-specific dynamics such as differences in translational kinetics, gene regulation, protein stability, and BBB permeability. This dissociation between *Mmp9* transcript levels and MMP-9 protein expression in the striatum suggests that peripheral sources and/or post-transcriptional mechanisms contribute to MMP-9 accumulation in the brain following Poly(I:C) exposure. However, further work is needed to clarify these mechanisms. For example, emerging evidence indicates sex-biased miRNA expression in the brains of alcoholics [42, 43]. The sustained elevation of MMP-9 in the male striatum from 6-24 hours can promote ongoing neuroplastic remodeling, which may influence neuronal function. It is worth noting that elevated striatal MMP-9 levels have been associated with BBB disruption [44], loss of dopaminergic neurons [45], alteration of reward circuitry [46], and alcohol dependence [47], positioning MMP-9 as a compelling target for further investigation in AUD-related neuroimmune mechanisms.

Strong mRNA–protein correlations for striatal CCL2, CCL5, and IL-6 suggest that TLR3-induced expression of these cytokines is primarily controlled by transcriptional regulation. These cytokines may play a prominent role in modulating striatum-dependent behaviors. In contrast, IL-1β and IFN-γ showed relatively weak correspondence between transcript and protein levels, suggesting the involvement of post-transcriptional regulatory mechanisms. Notably, for IL-1β and IFN-γ, a significant mRNA–protein correlation was observed in males but not in females, indicating a sex-dependent difference in this regulation. These findings suggest that post-transcriptional control may be more pronounced in females. The negative correlation between MMP-9 mRNA and protein levels in males re-emphasizes the high likelihood of externally derived MMP-9 (most likely from the blood) entering the striatum due to increased BBB permeability. Collectively, these findings demonstrate that neuroimmune responses are governed by multilayered regulatory mechanisms that vary by cytokine and sex, influencing post-transcriptional control and protein processing kinetics. Importantly, these data underscore that transcriptional changes alone do not reliably predict functional protein dynamics in the central nervous system following TLR3 activation, emphasizing the complexity of neuroimmune regulation.

Our findings show that TLR3 activation triggers coordinated, but sex-specific, immune signaling in both peripheral and central regions, leading to highly regulated time-dependent inflammatory responses that may affect behavior in AUD. Notably, MMP-9 activity seems to contribute to sex differences in alcohol-related behaviors. At the same time, CCL5’s expression in dopaminergic systems makes it a key mediator between immune activation and reward circuits. These results highlight the importance of temporal and tissue-specific immune regulation in shaping alcohol-related outcomes and emphasize sex as an essential biological variable. Overall, this research identifies potential molecular targets and supports the development of sex-specific immunomodulatory treatments aimed at preventing the progression from moderate to excessive alcohol consumption.

## Supporting information

https://osf.io/v47am/overview?view_only=1208a81f51204a57aa10976673b9b5bb

## Statements

### Data availability statement

The datasets presented in this article are not readily available due to privacy and ethical restrictions. The datasets are available upon reasonable request from the corresponding author.

### Ethics statement

All animal procedures were carried out in accordance with the guidelines of the National Institutes of Health (NIH) for the care and use of laboratory animals. The study protocol was reviewed and approved by the Institutional Animal Care and Use Committee (IACUC) at Texas Tech University Health Sciences Center. All efforts were made to minimize animal suffering and to reduce the number of animals used.

### Generative AI statement

The author(s) confirm that generative AI was not used to create the scientific content of this manuscript. However, limited AI-based tools were used solely to assist with language editing to improve clarity and coherence in selected sections.

## Supplementary material

The Supplementary Material for this article can be found online at:

## Author contributions

Philip Antwi-Adjei designed and performed the experiments, conducted data analysis, and wrote the manuscript. Shanmugam Sambantham contributed to experimental planning and assisted with performing experiments. Igor Ponomarev provided conceptual guidance, supervised the project, and contributed to manuscript writing, discussion development, and editing. Brent Kisby provided scientific advice and feedback throughout the study. All authors reviewed and approved the final manuscript

## Acknowledgements

We would like to thank Dr Scott Trasti and the staff of LARC/TTUHSC for their technical assistance with the breeding and animal experiments. Diagram schematics were generated using BioRender.

## Supplementary Figures

**Supplementary Figure 1:** Sex-specific expression of blood, striatum, and PFC *Ifnb1* 0, 6, 24, and 48 hours after PIC injection in FVB/B6 F1 hybrid mice.

**Supplementary Figure 2:** Sex-specific expression of immune proteins in striatum at 0, 6, 24, and 48 hours after PIC injection in FVB/B6 F1 hybrid mice.

