## Supplementary figures and images for "EFFECTS OF TOLL-LIKE RECEPTOR 3 – DEPENDENT IMMUNE ACTIVATION IN MICE ARE SEX- AND TISSUE- SPECIFIC: IMPLICATIONS FOR ALCOHOL USE DISORDER"

### https://osf.io/v47am/overview?view_only=1208a81f51204a57aa10976673b9b5bb

#### Supplementary Figure 1.

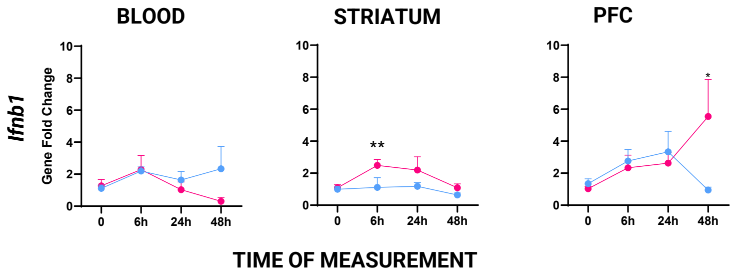


#### Supplementary Figure 2.

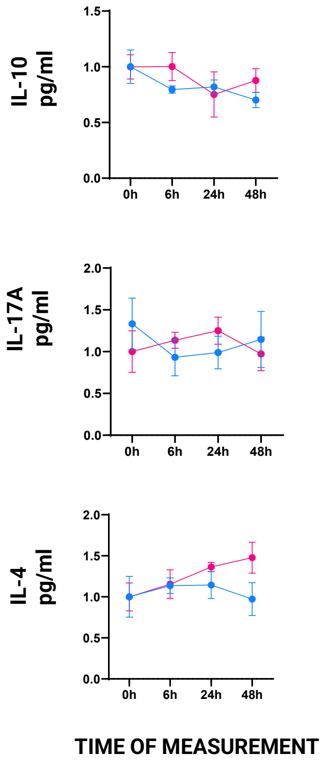
